# DBPOM: a comprehensive database of pharmaco-omics for cancer precision medicine

**DOI:** 10.1101/2021.01.18.427205

**Authors:** Yijun Liu, Fuhu Song, Zhi Li, Liang Chen, Ying Xu, Huiyan Sun, Yi Chang

## Abstract

During the course of cancer treatment, both efficacy and the adverse effects of drugs on patients should be taken into account. Although some public databases and modeling frameworks have been developed through studies on drug response, the negative effects of drugs are always neglected. Furthermore, most of them only considered the ramifications of the drug on the cell line, but the effects on the patient still require a huge amount of work to integrate data from various databases and calculations, especially in relation to precision treatment. In order to address these issues, we developed the DBPOM (http://www.dbpom.net/, a comprehensive database of pharmaco-omics for cancer precision medicine), which explores various drugs’ efficacy levels by calculating their potency in reverse, or enhancing cancer-associated gene expression changes. When compared with existing databases, the DBPOM could estimate the effectiveness of a drug on individual patients through the mapping of various cell lines to each person according to their genetic mutation similarities. The DBPOM is an easy-to-use and one-stop database for clinicians and drug researchers to search and analyze the overall effect of a drug or a drug combination on cancer patients as well as the biological functions that they target. We anticipate that DBPOM will become an important resource and analysis platform for drug development, drug mechanism studies and the discovery of new therapies.

## Introduction

Patients of the same cancer type, or even those at the same stage, may respond differently to the same drug[1, 2]; hence, a comprehensive study of drug screens and the mechanism of drug actions on patients with specific pathological and/or physiological conditions will be needed to facilitate improved individualized therapy[3, 4]. Pharmaco-omics is the use of genomics and transcriptomics-based information to help guide individualized drug therapy to maximize drug efficacy and minimize adverse drug reactions[5]. In the early stages of a drug’s development, its efficacy, toxicity, and sensitivity are typically tested on cell lines[6]. Currently, the potential treatment outcomes of a drug on a cancer might be assessed by using certain indices like half-maximal inhibitory concentration (IC50). In addition, another increasingly accepted measurement is the reversed expressions of genes, which are oppositely regulated between drug treatment information and disease-state data[7]. However, the adverse effects of drugs are usually ignored, which is a major concern for both public health and the development of medicines as failing to identify these negative outcomes could lead to significant amounts of morbidity[8]. Consequently, fully considering both reversed expressions and adverse effects is a highly desirable strategy[9, 10]. What is more, as cancer cells may develop drug resistance, it has been suggested that treatments that employ drug combinations potentially enhance efficacy and reduce toxicity[11]. Therefore, the mechanism of synergy and new combination recommendations have acted as a catalyst for intensive studies by academic researchers and pharmaceutical enterprises.

Recently, with the development of high-throughput technology, large amounts of cancer multi-omics data and related therapy data have been released that facilitate studies. The Cancer Genome Atlas (TCGA, https://www.cancer.gov/tcga)[12], molecularly characterizes over 20,000 primary cancer and matched normal samples, spanning 33 cancer types with detailed clinical information. Meanwhile, the Gene Expression Omnibus database (GEO, https://www.ncbi.nlm.nih.gov/geo/)[13] is a widely used public data repository of array and sequence-based multi-omics profiles, but it is not limited to cancer cells and samples. Also, the Cancer Cell Line Encyclopedia (CCLE, https://portals.broadinstitute.org/ccle)[14] collects and collates genomic, epigenomics data and transcriptomics data of more than 1000 cancer cell lines.

Some public databases and modeling frameworks have also been developed and widely used for studies of cancer precision medicine and drug response, such as CMap[15], GDSC[16], DSigDB[17], and CiDD[18]. While useful, most of these databases and computational tools only consider the effect of the drug on the cell line but not on cancer tissues, resulting in a huge amount of work to integrate data from various databases, and then calculation is still required to evaluate both the therapeutic and adverse effects on the patient, particularly during precision treatment, which makes it extremely difficult for clinicians and drug researchers to utilize them[19]. Here, we propose the creation of an open source, comprehensive database of pharmaco-omics for cancer precision medicine that is supported by the collection and storage of drug information, and analysis of the outcomes of drug integrative effect at both a cellular and individual patient level.

In order to evaluate the effect of a drug on individual patients and not just at a cellular level, we quantify both reversed and adverse outcomes by calculating differential expressions of all the genes between the cancer sample and the normal sample, the cancer cell line with and without drug treatment, respectively, and the mapping between cell lines and cancer tissues based on similarities in genetic mutations. By defining the score function through the presentation of the integrated drug effect and applying it to FDA-approved medicines, we find that most FDA-approved drugs, and the ones in clinical trials, tend to have a low DBPOM-defined score, which suggests that the DBPOM provides an effective means of appraising the usefulness and safety of drugs. It is also promising for precision medicine and drug-combination recommendations.

In summary, the DBPOM is an easy-to-use database for clinicians and drug researchers to search and analyze both the therapeutic and adverse effects of a drug/drug combination on cancer patients in accordance with the demand for personalized drug analysis, making it unlike similar databases that only focus on cell lines. By analyzing the biological function of reversed and adverse genes when different drug or drug combination treatments are being administered, the DBPOM can provide possible insights into studies of a drug’s mechanism and play a guiding role in drug development and recommendations. Furthermore, since the DBPOM provides a patient-centric module that focuses on the changes to the molecular features of each patient, it is naturally useful for querying and designing more effective personalized treatments.

## Methods

The overall design of the DBPOM is outlined in **Figure 1**.

**Figure 1.**
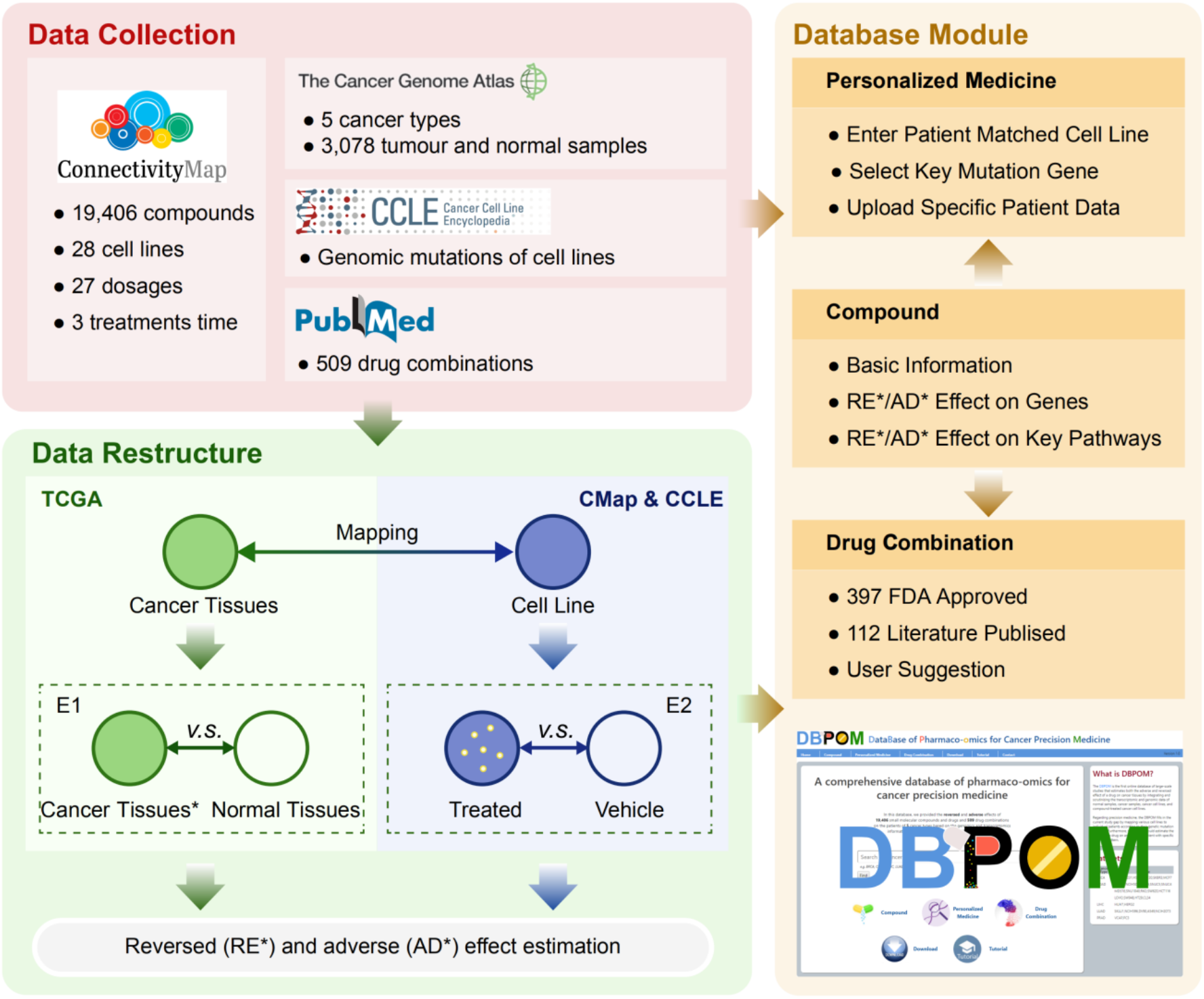
The procedures of the DBPOM database construction.

### Data source and pre-processing of the DBPOM

We collected genomics and transcriptomics data of over 3,000 cancer and normal samples across five cancer types from TCGA, and 28 relative cancer-cell lines from CCLE, respectively. This was as well as gathering more than 14,000 compound-perturbated gene expression profiles from CMap, nearly 4,000 IC50 profiles, which is a quantitative measure of drug efficacy from the GDSC, and over 500 examples of drug combination information from the literature[20]. In addition, five cancer types that each has at least two forms of cell line have been used for precision medicine analyses. In total, 28 cell lines and 3,078 cancer samples have been selected from the largest intersection between the CMap, the CCLE and TCGA. After removing 18 pseudogenes, we retained 12,310 expressed genes and their expression under various conditions from CMap. 19,406 small molecule compounds and drugs were determined via the selected 28 cell lines, 27 dosages and three treatment durations. We manually annotated the Ensembl ID of the gene in TCGA and the gene name from the CMap. Detailed information can be downloaded in the download module. All data was collected before September 2019. **Table 1** summarizes the key information stored in the database.

**Table 1.**
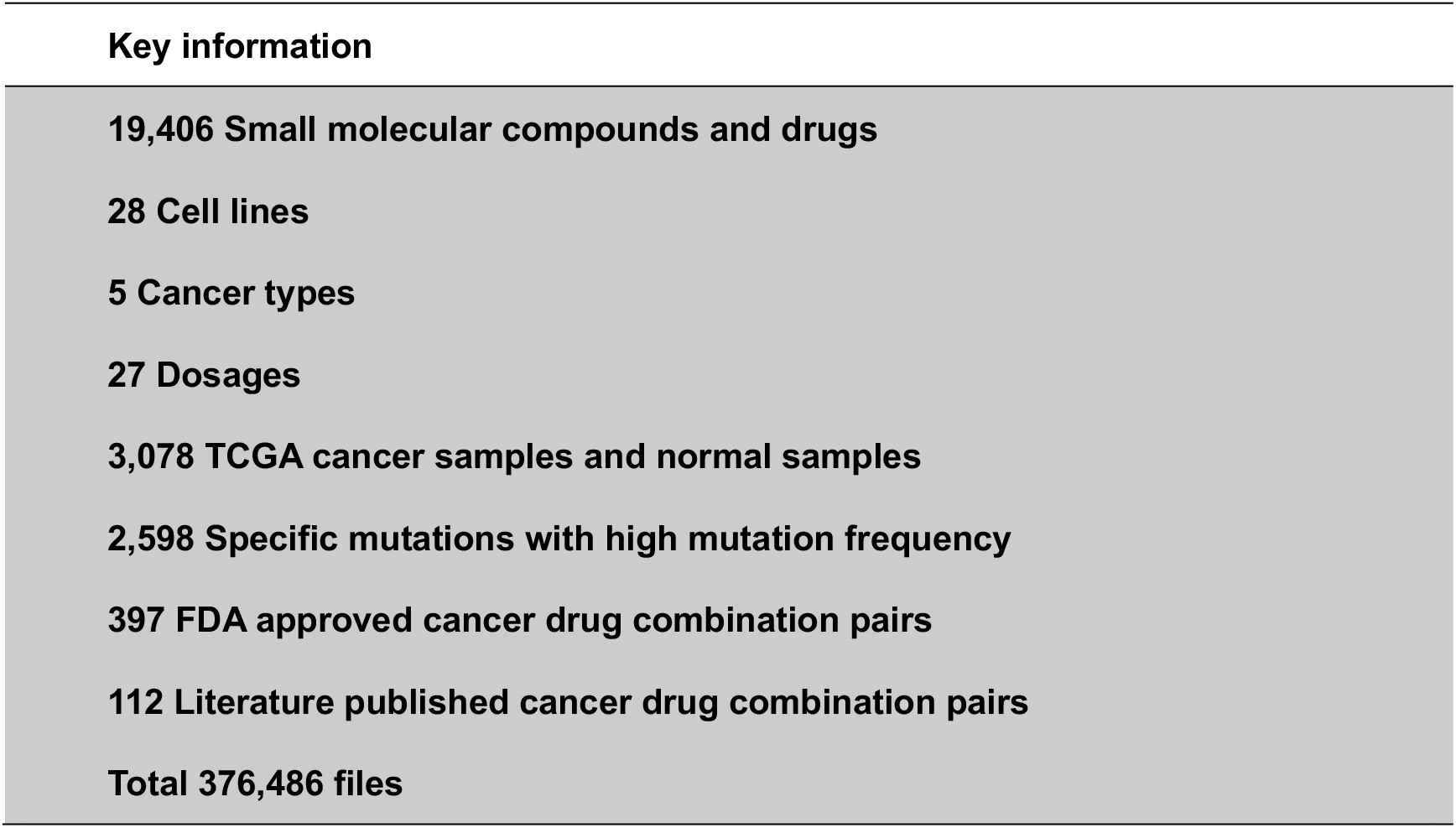
Statistical information about the DBPOM.

### Differential gene expression analysis and comparisons

Differential gene expression analyses have been performed on both cancer versus control tissues samples (E1) from TCGA and molecular-compound-treated cell lines versus DMSO-treated cell lines (E2) from the CMap database.

With regards to E1, to identify which genes are associated with carcinogenesis at the gene expression level, a DESeq2 package[21] was used for differential expression analysis between normal and cancer samples in TCGA. We set the threshold as p-value=0.001 and fold-change=2. For each cancer type, we defined the gene as up-regulated (down-regulated) and labelled it as 1 (−1) if its log2 fold-change is above 1 or less than −1 and the p-value is less than 0.001 when comparing cancer samples with normal samples; otherwise, it was labelled as 0.

In relation to E2, to evaluate the effect of compounds on the cell line at the gene level, the following formula was used for differential expression analysis between DMSO-treated and compound-treated cell lines in the CMap. We defined genes as significantly differentially expressed if their fold-change is above 15. For each cell line, we define the gene as up-regulated (down-regulated) and label it 1 (−1) if its fold-change is above 15 or less than −15 when comparing compounds with DMSO-treated cell lines; otherwise, it was labelled as 0.

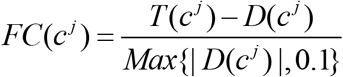

Where 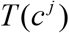 and 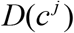 represent the mRNA expression value of the *j*-th gene in the compound treated group and the DMSO-treated group, respectively. 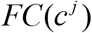 represents the fold-change value. In order to prevent overflow, we set 0.1 as the lowest expression value.

### Disease gene expression signatures and assessment

Within the DBPOM, we have developed a tool for distinguishing the reversed and adverse genes of a specified compound, or drug on cancer samples, by comparing the data stemming from the two groups of differential gene expression results (E1 and E2). When a cell line is treated with a compound, if the effect on a gene is oppositional, meaning it is up-regulated in E1 and down-regulated in E2 or vice versa, we consider it to have a reversed effect on said gene. Otherwise, if a gene is marked as 1(E1) ~1(E2), −1(E1) ~-1(E2), 0(E1) ~1(E2) or 0(E1) ~-1(E2), with 0 standing for no change, we consider the compound to have an adverse effect on this specific gene. For each compound, if it reverses the expression of a cancer-associated gene, it is thought to be effective on the gene. On the contrary, if the gene expression substantially deviates after treatment with the compound, we believe that the drug has had an adverse effect on the gene. It should be noted that the adverse effects defined here are different from the pharmacological adverse reactions in clinical trials.

The formulaic expression is as follows:

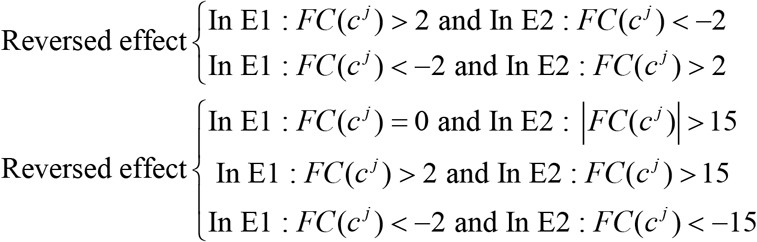

In order to estimate the overall effect of a drug on each patient, we integrated both the reversed and adverse effects as described above of all of the genes. Moreover, we defined the score as the reversed and adverse of drug effect (RADE) to indicate the safety and effectiveness of each compound. RADE comprehensively considered the number of reversed genes and adverse genes after being treated by the compound. The lower the score, the higher the effectiveness and safety. The formulaic expression is as follows:

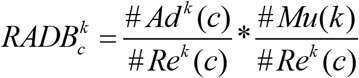

The need to explain pound sign (*#*) represents the number of elements in the set. In the above function, 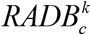 signifies the efficacy score of compound treatment on cell line *k*. 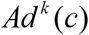 and 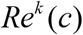 are the set of adverse and reversed genes in cell line *k* with the treatment of the compound, respectively. *Mu*(*k*) is the gene set that should be reversed according to previous TCGA analyses.

### Mapping individual mutation phenotype to cancer cell line

Genetic mutations of different cancer cell lines of the same cancer type vary, which results in them responding divergently to the same drug[22]. Meanwhile, as an important reference for molecular subtypes, genomic mutation could reflect the individual characteristics of different patients with the same type of cancer. Therefore, through the estimation of the cell line’s response to drugs and by establishing the association between cancer cell lines and the similarities between cancer patient-based genetic mutations, it is helpful for recommending personalized medicine.

We collected the genomic mutation information of each cell line and each cancer patients from the CCLE and TCGA database, respectively, and defined the similarity score of each cancer cell line and cancer tissue of the same cancer type in the following manner:

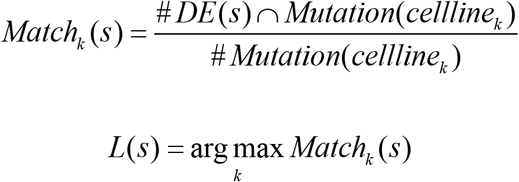

*Match*_*k*_(*s*) is the matching score of the sample on cell line *k* and the sample is from corresponding cancer type provided by TCGA. *DE*(*s*) is the set of mutation genes in the sample and *Mutation*(*cellline*_*k*_) is the set of mutation genes of cell line *k*. *L*(*s*) represents the sample from TCGA, which is roughly labelled as a cancer type in TCGA, and it has been chosen for re-labelling as the cell line with the highest matching score.

There are three reference options for mapping according to the samples selected from E1 and the mapping method we proposed:

i. TCGA versus CMap: in E1, all of the cancer samples of the selected cancer type in TCGA are utilized. While this is the general strategy in current research, it does not consider individual differences. We provide this option in case some users need it.
ii. subTCGA-mutaion versus CMap: in E1, the subset of cancer samples in TCGA which have high matching scores with selected cell line are utilized. We provide this strategy as overall mutation pattern is an important criterion for distinguishing different cell lines. So, we provide users with this option to suit diverse requirements. In practice, for each cancer patient, he or she is firstly classified into the most suitable sample group according to mutation similarity, and then to analyze effect of drugs on him or her.
iii. subTCGA-specific versus CMap: E1 utilizes a subset of cancer samples with the specific mutation that user concerns from TCGA, which match the specific mutation that user concerns of the selected cell line in E2. We proposed this strategy and expected to get a reasonable evaluation of drug effectiveness that could then assist precision medicine.

### Pathway enrichment analyses

To better disclose the real effects of drugs on cancer samples, we performed function analysis on the adverse and reversed genes of each perturbation experiment to find the main biological pathways that the compound potentially targets. Additionally, the DBPOM provides special enrichment analysis of 11 key pathways that concern researchers the most, which are closely related to cell proliferation, death and apoptosis. The gene sets of all the biological processes and pathways were collected from GSEA[23], and the statistical significance of the enrichment analyses was obtained based on the hypergeometric test below:

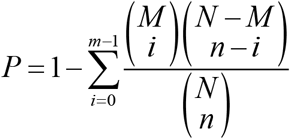

Here, *N* is the number of all genes in the transcriptome, and *M* is the number of genes in the gene set to be detected. Meanwhile, *n* is the number of genes intersecting with the adverse/reversed gene set in *N*, *m* is the number of adverse/reversed genes in the pathway, and *P* represents the enrichment significance of adverse/reversed genes on the pathway.

## Results

The DBPOM is designed to evaluate the effect of treatment when using each compound on every cancer patient by integrating the gene signatures of different perturbation experiments on cell lines, particularly cancer types or subtypes associated with genes. The database contains three modules, which are the compound effect on cancer samples, patient-centric precision medicine and drug effect comparison of known/unknown drug combination pairs.

### Search by compound

This module is compound-centric and contains response information of each compound at different dosages and treatment lengths when applied to all cell lines. As shown in **Figure 2.A**, when a user selects a compound, the response information has two levels. One is the whole cell level. Users can bring up the detail page for a more in-depth understanding and pathway enrichment analyses of KEGG, GO, PID and REACTOME, which are provided for the purpose of analyzing the functions of reversed and adverse genes. The other stratum is the pathway level. The database provides a panel focusing on the treatment effects of drugs or compounds on the genes involved in cell proliferation, cell death and 11 other key pathways of high attention for clinicians and drug researchers, including KEGG_CELL_CYCLE and apoptosis regulation. The module also offers information about the compound like its IC50 and structure from other databases. Users can choose a cell line in isolation to ascertain the responses of all compounds to this cell line at different dosages and treatment durations. According to our calculation formula, a drug with a higher score is thought to have a lower possibility of being effective on a certain cell line.

**Figure 2.**
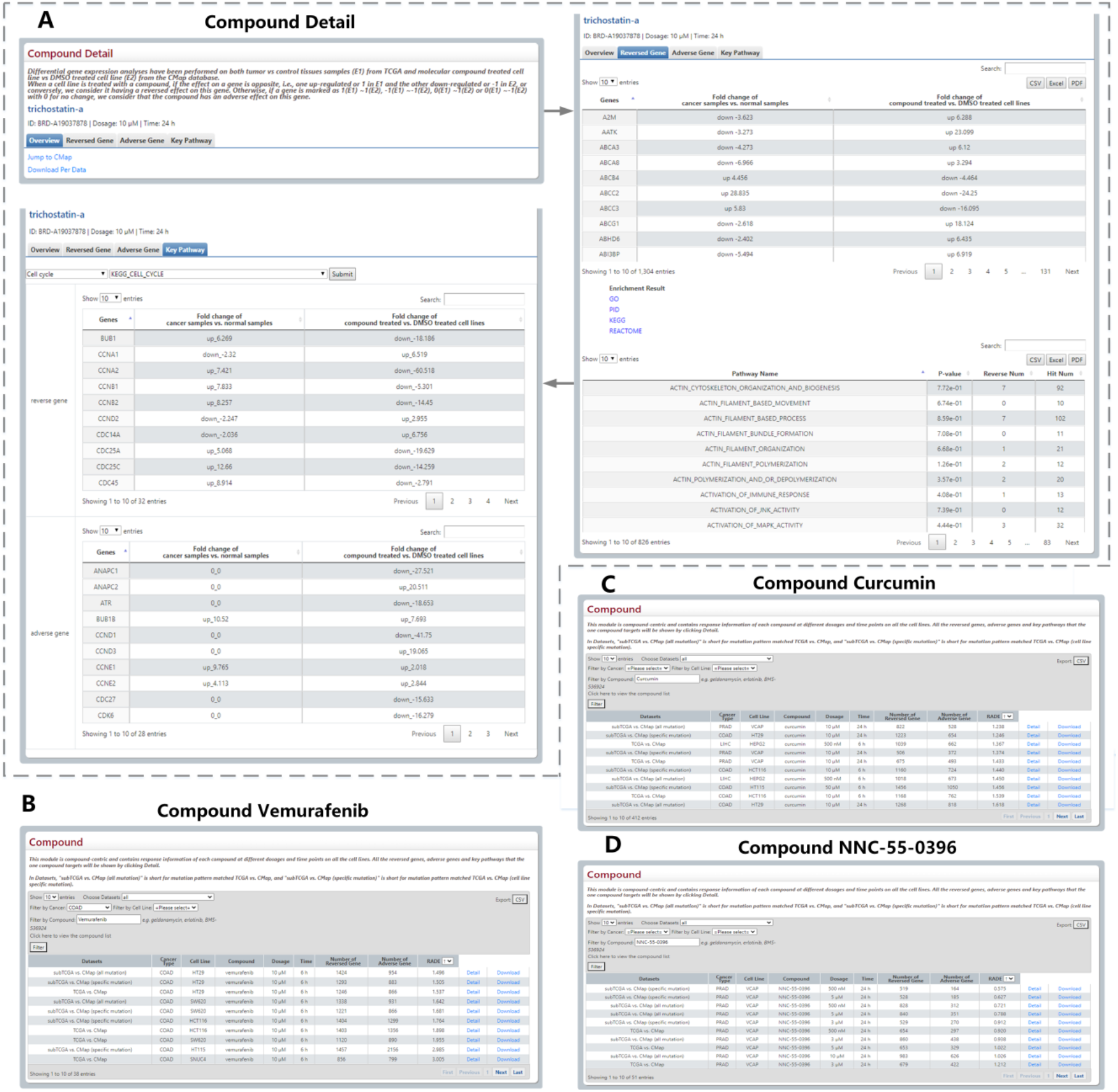
Outline and statistical analysis of Compound module. (A) The compound detail page includes concise information, details of reversed and adverse genes, and pathway enrichment analysis. (B–D) Three responsive tables of users’ search results have been employed to prove the rationality of the RADE scores raised by the DBPOM.

We found that several drugs that were FDA approved, and some that are being employed in clinical trials for cancer treatment, have low RADE values in relation to our database, which demonstrates that the score is reasonably successful at evaluating the drug’s efficiency. For example, vemurafenib is an FDA-approved drug used for the treatment of metastatic melanoma with a mutation on the BRAF in the valine located in the exon 15 at codon 600[24] When combined with standard-of-care or novel-targeted therapies, it is also reported that vemurafenib is effective on colorectal cancer[25]. Via our database, we discovered that the RADE value of vemurafenib acting on COAD generally ranks extremely highly as well (**Figure 2.B**). As another example, curcumin is a drug that has been used in clinical trials, having been investigated for the treatment and supportive care of clinical conditions, including proteinuria, breast cancer, multiple myeloma, and non-small-cell lung cancer[26]. The RADE value of curcumin ranks very highly on the whole in our database (**Figure 2.C**). Moreover, NNC-55-0396 is a molecular compound still in the drug development phase and reportedly could prevent human cancer cell proliferation and induce cancer cell apoptosis as a result of its ability to inhibit the function of T-type Ca^2+^ channels[27]; the experimental results are also in good agreement with the value we proposed (**Figure 2.D**).

### Personalized medicine

This is a patient-centric module and contains three options for precision medicine. The first is for an optimal match between a cell line and cancer samples according to the similarity score of their overall mutation patterns. When a cell line is chosen, the module matches the referenced cancer samples with the cell, and performs the aforementioned analyses (**Figure 3.A**). The second example relates to the specific mutation that a user is interested in. When a certain mutation and a drug are selected, this module compares both the reversed and adverse effects of this drug on cancer patients versus one without the mutation (**Figure 3.B**). For each cancer, we only list the genes which have high genetic mutation rate. Finally, the user can also submit his/her own data regarding a patient’s mutation profile for analysis (**Figure 3.C**).

**Figure 3.**
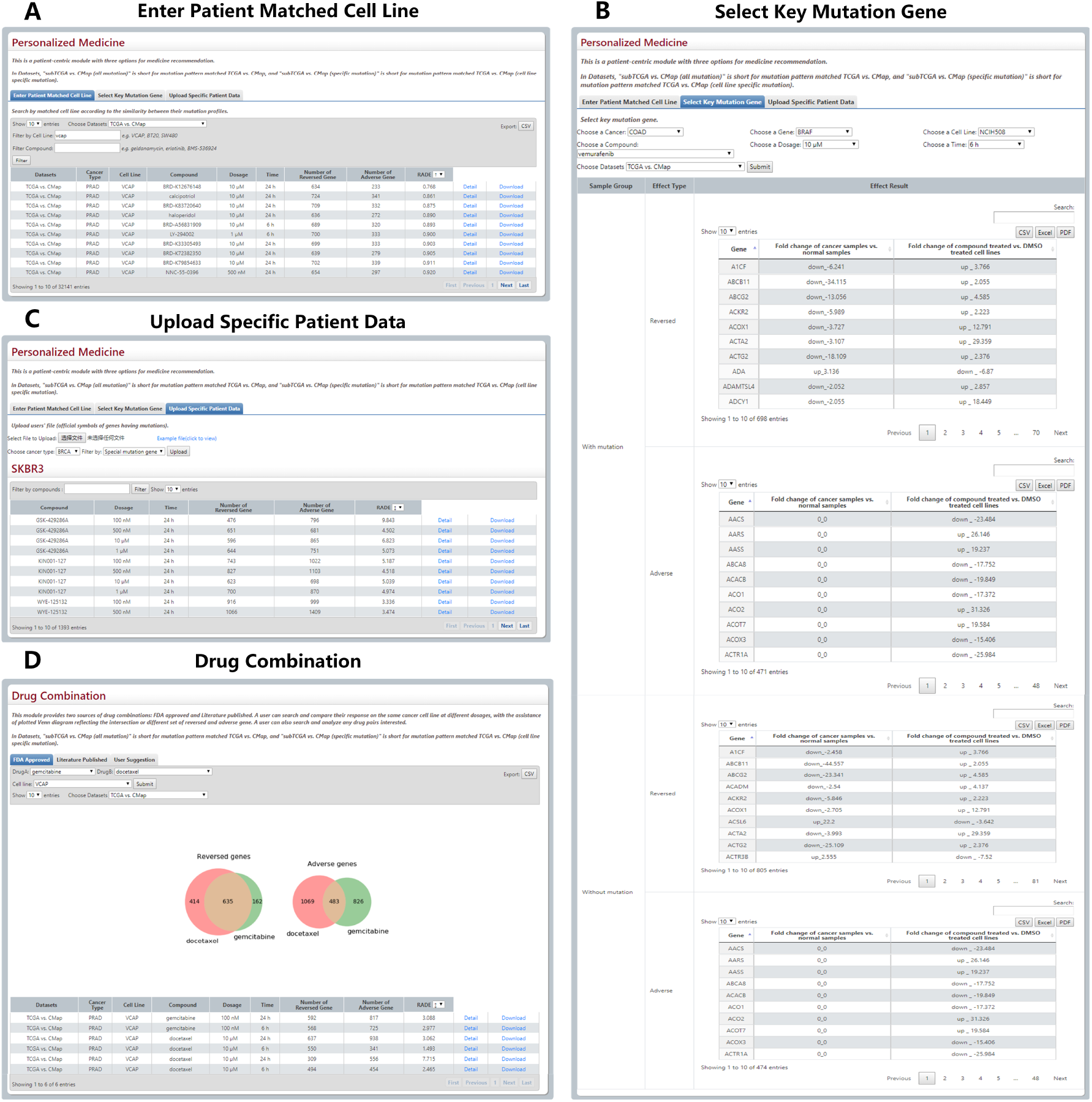
Results from Personalized Medicine and Drug Combination modules. (A–C)The three modules for personalized medicine. (D) The Venn plot and search results of gemcitabine and docetaxel.

### Drug combination

This module provides FDA-approved, text-mining-validated drug pairs. For each pair, the user can search and compare their response on the same cancer cell line in a different physical environment with the assistance of a plotted Venn diagram that reflects the intersection or a different set of reversed and adverse genes. Furthermore, they can also search and investigate any drug pairs they are interested in. For instance, gemcitabine and docetaxel are an FDA-approved drug combination used for breast, ovarian, and non-small-cell lung cancer. Their reversed and adverse genes on VCAP cell line and the corresponding Venn plot are shown in **Figure 3.D**.

### Easy Download

This module allows easy access to all the reversed and adverse effects of 19,406 small molecular compounds, existing drugs and 509 drug combinations on the 12,310 genes of cancer patients across five cancer types based on 28 cell lines and 3,078 cancer samples. In addition, all of the documents used in the database are available for download, which is also the case with the compounds, key pathways and drug combinations.

### Database and web interface service and implementation

The DBPOM website displays succinct information about the gene signatures of compounds and drugs. Detailed information can also be downloaded from the relevant pages. We provide partial/complete downloads of the entire database and its description files. Furthermore, the DBPOM website affords researchers an online analysis capability for compounds, and they can easily download their query or calculation results from any module. The data in the DBPOM is all stored and managed by MySQL, and the web interface is implemented using HTML and JavaScript. In addition, all of the data analysis work is completed with R and Python.

## Discussion and future directions

Overall, DBPOM is an easy-to-use database, which offers users the chance to explore the mechanism of the effect of a potential drug on cancer patients via the use of genomics and transcriptomics-based information. Clinicians and drug researchers could search and analyze both the therapeutic and adverse effects of a drug or drug combination on cancer patients according to their needs for personalized drug analysis, which is unlike other similar databases that focus solely on cell lines. Among other advantages, the DBPOM provides a worthwhile way to compare the effect of different drugs through the prism of gene expression change, meaning it could assist in speeding up the process of drug development, facilitate new uses of old drugs, and act as a catalyst for the discovery of new therapies. In addition, the DBPOM is the first large-scale database to study adverse gene expression during drug treatment.

We will continue to collect more data relating to other cancer types, cell lines, compounds, and approved drug combinations to enrich our database. We will also develop and import more computational tools for reversed effects, adverse effects and their comprehensive effect estimation as well as extending the capabilities for precision medicine recommendations. We expect the DBPOM to become an important resource and analysis platform for drug development, drug mechanism studies, and new therapy discoveries.

## Acknowledgments

The authors thank all members of the Zhi Li Laboratory at the First Hospital of China Medical University who provided us with critical comments and feedback.

## Funding

National Natural Science Foundation of China (61902144, 62002212, 62072212) and 2020 Li Ka Shing Foundation Cross-Disciplinary Research Grant (2020LKSFG07D).

## Notes

### Competing Interest Statement

The authors have declared no competing interest.

